# Overlap of competing voices may mask attentional effects on speech-like distortion product otoacoustic emissions

**DOI:** 10.64898/2026.11.11.765909

**Authors:** Janna Steinebach, Tobias Reichenbach

**Affiliations:** Department Artificial Intelligence in Biomedical Engineering, Friedrich-Alexander-Universität Erlangen-Nürnberg, Nürnberger Strasse 74, 91052 Erlangen, Germany

**Keywords:** otoacoustic emissions | selective attention | medial olivocochlear system | efferent feedback | speech | distortion products | replication

## Abstract

Whether selective auditory attention modulates the cochlear amplifier through the medial olivocochlear system remains debated. Using speech-like distortion-product otoacoustic emissions (DPOAEs), elicited by waveforms that follow the harmonics of natural speech, we previously found that selectively attending one of two competing talkers reduced the emissions related to the resolved harmonics of the female voice. Here, we attempted to reproduce this effect in a design that separated different frequency regimes and harmonic resolvability. Forty-four participants at-tended one of two competing talkers, a female and a male voice, presented to one ear, while speech-like DPOAEs related to both voices were recorded from the other ear. Stimuli were derived from resolved and unresolved harmonics in a low-frequency (up to 1.5 kHz) and a transitional (1.5–4 kHz) regime, which required pitch-shifting both voices so that their fundamental frequencies were much closer to each other than in the original study. Speech-like DPOAEs emerged for most stimulus types, although they were weak for some and absent for one. Their amplitude, however, did not depend on attention for any stimulus type. Further time-domain metrics showed only isolated attentional effects of opposite direction for two stimulus types, and a frequency-domain metric showed none. A cochlear excitation model suggests that, due to the closely spaced fundamental frequencies of the two voices, the emissions most comparable to the original study presumably arose at cochlear locations shared by both voices, where attention would modulate both voices alike.

## 1 INTRODUCTION

Humans excel at following one talker among several, an ability known as the cocktail party effect [1]. The neural basis of this ability has mostly been studied in the auditory cortex. However, the auditory system also contains extensive descending pathways, which reach down to the brainstem and the cochlea [2, 3]. Their final stage is the medial olivo-cochlear (MOC) system, whose fibres innervate the outer hair cells and can reduce the gain of the cochlear amplifier [4]. Because the MOC innervation follows the tonotopic map of the cochlea, this gain could in principle be adjusted in a frequency-specific manner, for instance to favour the frequencies of an attended talker. Such frequency-specific gain control has been proposed to aid speech perception in noise [5, 6].

Otoacoustic emissions (OAEs) offer a non-invasive window onto the cochlear amplifier and have therefore been used to test whether attention modulates cochlear activity. The results have been inconsistent. Some studies reported reduced or enhanced emissions during attention [7–10], others found no effect [11], and attempts to replicate reported effects were partly unsuccessful [12–15]. One reason may be that most studies elicited the emissions with clicks or pure tones, which differ markedly from the speech signals for which selective attention is most relevant.

Speech-like distortion-product OAEs (DPOAEs) were developed to overcome this limitation [16]. They are elicited by two waveforms that follow two harmonics of the voiced parts of a speech signal, so that the emission reflects cochlear activity at the frequencies of that speech. Using one stimulus pair per voice, Saiz-Alía et al. [16] found that the speech-like DPOAEs related to the female voice were larger when this voice was attended than when it was ignored. In a subsequent study, which we refer to as JS25, we presented four stimulus pairs at the same time, derived from resolved and unresolved harmonics of a female and a male voice [17]. Attention strongly reduced the speech-like DPOAEs related to the resolved harmonics of the female voice: their amplitude was about 20 % smaller when this voice was attended than when it was ignored (*p* = 0.0003). The emissions related to the unresolved harmonics of the male voice were modulated as well, which we attributed to their spectral overlap with the resolved harmonics of the female voice. JS25 nevertheless left three questions open. First, the emissions related to the resolved harmonics of the male voice could not be measured reliably because of their low frequencies, so that the counterpart to the effect for the female voice remained untested. Second, the spectral overlap between the stimuli of the two voices made it difficult to attribute an effect to one voice. Third, the direction of the effect was unexpected and opposite to that observed by Saiz-Alía et al. [16]: the emission was smaller, not larger, when the corresponding voice was attended. We hypothesised that this direction is specific to the low-frequency, apical region of the cochlea, whose mechanics differ from those of the base [17, 18], so that attention might reduce the emissions in apical regions but enhance them in basal ones. A simpler explanation would be the inhibitory action of the MOC system itself. MOC activation reduces the amplification by the outer hair cells and thereby makes the compressive input–output function of the basilar membrane more linear [4, 7, 19], which weakens the emissions that arise from this nonlinearity. Such a linearisation has been proposed to aid speech perception in noise by attenuating the background evenly while keeping the structure of speech salient [7]. If attention increased the MOC activity at the places excited by the attended voice, it would thus reduce the emissions related to this voice in any region of the cochlea, not only in apical ones.

The present study, JS26, was designed to address these questions. We separated two factors that were confounded in JS25: the frequency region of the stimuli, and whether they were derived from resolved or unresolved harmonics. To this end, we presented only one stimulus pair per voice at a time, avoided spectral overlap between the stimuli of the two voices, and shifted the fundamental frequencies of both voices so that their stimuli fell into either a low-frequency or a transitional frequency regime. We expected to reproduce the effect of JS25 for the resolved harmonics in the low-frequency regime, now for both voices, and to test whether its direction reverses at higher frequencies.

Contrary to this expectation, attention did not robustly modulate the speech-like DPOAEs. We report this result together with an analysis of whether the study was sensitive enough to detect an effect of the size observed in JS25. We also discuss which differences in the experimental design might explain the different outcomes.

## 2 MATERIALS AND METHODS

### 2.1 Participants

Forty-four participants took part in the study (22 female, 22 male; aged 20–43 years, mean *±* SD: 27.3 *±* 3.4 years). All were right-handed native speakers of German and reported no neurological disorders or hearing impairments. No participant was excluded. The study was approved by the ethics committee of the University Hospital Erlangen (registration 133-12B), and all participants gave written informed consent.

### 2.2 Overview of the paradigm

The paradigm followed that of JS25 [17] (Fig. 1). Two audiobooks, one read by a female and one by a male narrator, were mixed and presented to the right ear. Participants attended either the female or the male voice and ignored the other. At the same time, two speech-like DPOAEs, one related to each voice, were elicited and recorded in the left ear. Because both emissions were recorded in the same trial, the attended state of one voice was always recorded together with the ignored state of the other, and the acoustic input was identical in both attentional conditions. We refer to the attentional conditions as attended (Att.) and ignored (Ign.), always with respect to the voice from which a stimulus was derived.

**Figure 1:**
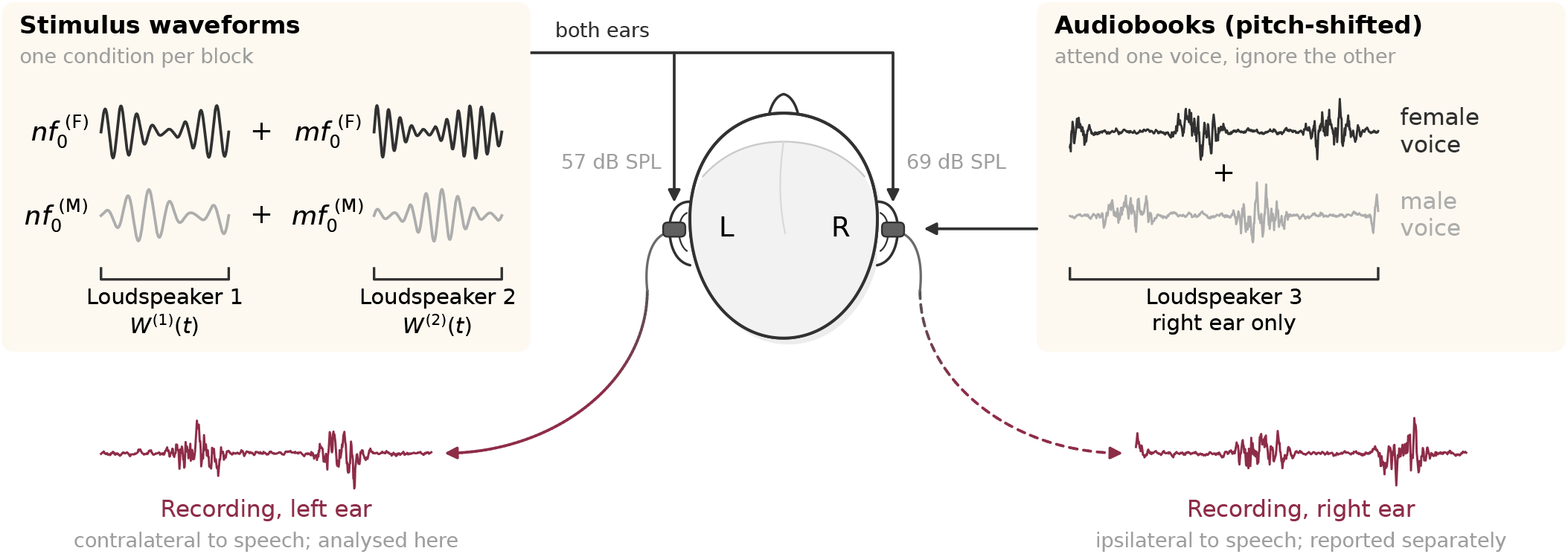
Experimental setup. Two audiobooks, read by a female and a male narrator and pitch-shifted separately for each condition, were mixed and presented to the right ear through one loudspeaker of the probe (right). Participants attended one voice and ignored the other. In each block, one stimulus pair per voice was derived from the harmonics *n* and *m* of its fundamental frequency *f*_0_ (left). The two waveforms with harmonic number *n* were summed to form *W* ^(1)(^*t*), and the two with harmonic number *m* to form *W* ^(2)(^*t*). Both sums were delivered to both ears through two further loudspeakers. Average sound pressure levels were 57 dB SPL in the left ear (stimulus waveforms) and 69 dB SPL in the right ear (audiobooks and stimulus waveforms). Recordings were obtained from both ears; the present study analyses only the recordings from the left ear, contralateral to the speech.

Compared with JS25, we changed the design in eight respects. Five changes concern the stimulus design and are likely to have affected the outcome: 1) only two stimulus pairs instead of four were presented at a time; 2) the stimuli were placed in predefined frequency regimes; 3) the fundamental frequencies of the voices were shifted to achieve this; 4) the filter that determines the bandwidth of the stimuli was narrower; and (5) the audiobooks were presented at a higher sound-pressure level. Three further changes are minor: 6) natural instead of synthesised voices were used; 7) the stimulus waveforms were presented to both ears, to allow additional recordings ipsilateral to the speech, which are not analysed here; and 8) each condition was repeated three instead of six times, to keep the total duration of the experiment within 60 minutes.

### 2.3 Speech material and pitch shifting

The speech material consisted of two commercial audiobooks: “Den Hund überleben” [20], read by a male narrator, and “Eine Frau erlebt die Polarnacht” [21], read by a female narrator. Unlike in JS25, we used natural rather than synthesised voices, because they sounded considerably more pleasant after pitch shifting. The two audiobooks were scaled so that they produced the same sound pressure level, averaged over two-minute segments, in test recordings both in the ear and outside of it.

To place the stimulus pairs of both voices in the same frequency regime, the fundamental frequencies of the voices had to be shifted, separately for each condition. We used pitch-synchronous overlap-add (PSOLA) resynthesis in Praat [22], accessed through Parselmouth [23]. The fundamental frequency contour *f*_0_(*t*) of each segment was shifted on a logarithmic scale so that its median matched the target value, which preserved its relative variability. Three iterations of resynthesis and re-estimation of the median *f*_0 c_orrected residual deviations from the target.

The resulting fundamental frequencies are listed in Table 1. They were chosen under four constraints: the stimulus waveforms of both voices had to fall into the intended frequency regime; the two voices had to be separated by the same amount in all conditions; the shifted voices had to remain intelligible and pleasant enough to be followed for two minutes; and the correlation between the stimulus waveforms and the waveform of the expected emission had to remain small. We quantified the separation of the two voices as the difference of their mean fundamental frequencies on the ERB-number scale [24],

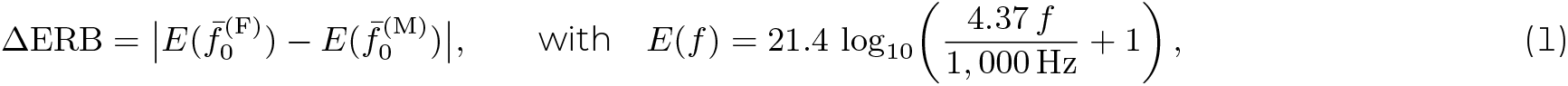

in which *f̄*^(F)^_0_ and *f̄*^(M)^_0_ are the mean fundamental frequencies of the female and the male voice. ΔERB thus gives the distance between the fundamental frequencies of the two voices in auditory filter bandwidths. Because both voices had to fit into the same frequency regime, ΔERB was only 0.9 to 1.0, compared with 2.6 in JS25 and 1.7 in the study of Saiz-Alía et al. [16].

**Table 1:**
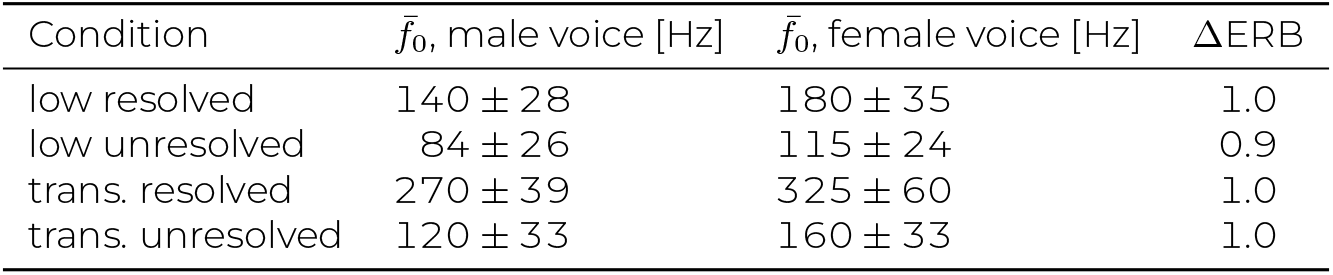
Mean fundamental frequencies *f̄*^0^ of the pitch-shifted voices in the four conditions (mean *±* SD across all two-minute segments) and the resulting separation of the voices (Eq. 1).

| Condition | f̄<sub>0</sub>, male voice [Hz] | f̄<sub>0</sub>, female voice [Hz] | ΔERB |
|---|---|---|---|
| low resolved | 140 ± 28 | 180 ± 35 | 1.0 |
| low unresolved | 84 ± 26 | 115 ± 24 | 0.9 |
| trans. resolved | 270 ± 39 | 325 ± 60 | 1.0 |
| trans. unresolved | 120 ± 33 | 160 ± 33 | 1.0 |

### 2.4 Stimuli for eliciting speech-like DPOAEs

The stimuli were generated as in previous work [16, 17, 25]. In brief, the voiced parts of each pitch-shifted speech signal were bandpass filtered around *f̄*^0^ to obtain a fundamental waveform *w*_0(_*t*), which was normalised by z-scoring. Its analytic representation *W*_0(_*t*) = *w*_0_(*t*) + *i H*[*w*_0_](*t*) = *A*(*t*) *e^iϕ^*^(^*^t^*^),^ with *H* denoting the Hilbert transform, yields the instantaneous amplitude *A*(*t*) and phase *ϕ*(*t*). A waveform that follows the *n*-th harmonic of the voice is then given by

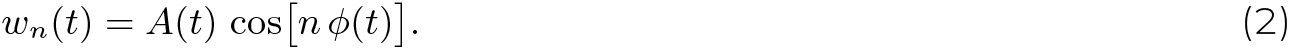

Two such waveforms *w_n_*(*t*) and *w_m_*(*t*) with *n < m* form a stimulus pair. Their cubic distortion product follows the harmonic 2*n − m* of the voice, so that the expected emission is described by the waveform *w*_2_*_n−m_*_(_*t*), which we refer to as the DPOAE template.

In JS25, the bandpass filter for *w*_0_(*t*) spanned *±*0.5 standard deviations of *f*_0_(*t*) around *f̄*^0^. Since the voices were much closer here, this filter would have caused the stimulus waveforms of one voice to correlate more strongly with the DPOAE templates of the other voice. To reduce these correlations and thereby improve the signal-to-noise ratio of the measurement, we used a fixed, narrow filter of *f̄*^0^ *±* 5 Hz for *f̄*^0^ *<* 100 Hz and *f̄*^0^ *±* 10 Hz otherwise. As a consequence, the stimuli and templates were considerably more narrowband than in JS25 and followed the slow variations of the pitch contour only within a few tens of hertz (cf. Table 2).

**Table 2:**
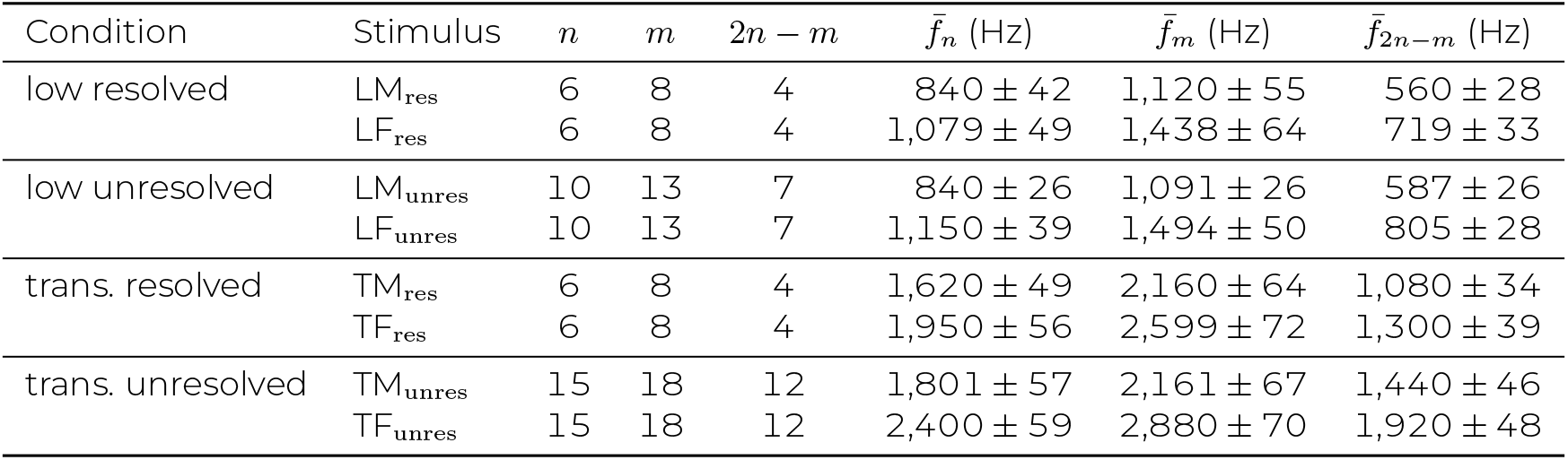
Harmonic numbers *n* and *m* of the stimulus pairs, harmonic number 2*n − m* of the DPOAE template, and mean frequencies *f̄ ^n^* = *nf̄*^0^(*t*), *f̄^m^* = *mf̄*^0^(*t*) and *f̄*^2^*n−m* = (2*n − m*)*f̄*^0^(*t*) of the waveforms (mean *±* SD across all two-minute segments), for the eight stimulus types. Horizontal lines separate the four conditions.

| Condition | Stimulus | *n* | *m* | 2*n* − *m* | $\bar{f}_n$ (Hz) | $\bar{f}_m$ (Hz) | $\bar{f}_{2n-m}$ (Hz) |
|---|---|---|---|---|---|---|---|
| low resolved | LM<sub>res</sub> | 6 | 8 | 4 | 840 ± 42 | 1,120 ± 55 | 560 ± 28 |
|  | LF<sub>res</sub> | 6 | 8 | 4 | 1,079 ± 49 | 1,438 ± 64 | 719 ± 33 |
| low unresolved | LM<sub>unres</sub> | 10 | 13 | 7 | 840 ± 26 | 1,091 ± 26 | 587 ± 26 |
|  | LF<sub>unres</sub> | 10 | 13 | 7 | 1,150 ± 39 | 1,494 ± 50 | 805 ± 28 |
| trans. resolved | TM<sub>res</sub> | 6 | 8 | 4 | 1,620 ± 49 | 2,160 ± 64 | 1,080 ± 34 |
|  | TF<sub>res</sub> | 6 | 8 | 4 | 1,950 ± 56 | 2,599 ± 72 | 1,300 ± 39 |
| trans. unresolved | TM<sub>unres</sub> | 15 | 18 | 12 | 1,801 ± 57 | 2,161 ± 67 | 1,440 ± 46 |
|  | TF<sub>unres</sub> | 15 | 18 | 12 | 2,400 ± 59 | 2,880 ± 70 | 1,920 ± 48 |

As in previous work, we distinguished between stimuli derived from resolved and from unresolved harmonics. Harmonics up to about the tenth are considered to be resolved by the cochlea, that is, they produce separate peaks along the basilar membrane, whereas higher harmonics overlap [26, 27]. As in JS25, the terms refer to the harmonics of the speech signal from which a stimulus pair was derived, not to the two waveforms of the pair, which have a similar frequency ratio *m/n* in all conditions. In addition, we distinguished between two frequency regimes of the stimulus waveforms: a low-frequency regime up to 1.5 kHz and a transitional regime between 1.5 and 4 kHz. Cochlear mechanics and tuning differ between apical and basal regions, with transitions reported around 1.5 to 2 kHz [18, 28, 29] and around 4 to 5 kHz [30–32], so that the transitional regime lies between these two transitions. A high-frequency regime above 4 kHz could not be included, because the required fundamental frequencies could not be reached by pitch shifting while keeping the speech intelligible.

Crossing the two factors resulted in four conditions: low resolved, low unresolved, transitional resolved and transitional unresolved. In each condition, one stimulus pair was derived from each voice, which yields eight stimulus types. Their names combine the frequency regime (L: low, T: transitional), the voice (F: female, M: male) and the harmonics from which they were derived (res: resolved, unres: unresolved): LF_res_ and LMres in the low resolved condition, LFunres and LM_unres_ in the low unresolved condition, TF_res_ and TM_res_ in the transitional resolved condition, and TF_unres_ and TM_unres_ in the transitional unresolved condition.

The harmonic numbers and the resulting frequencies are given in Table 2. The stimulus waveforms were placed in the respective frequency regime; the DPOAE templates necessarily lie at lower frequencies. Within these constraints, we kept the ratio *m/n* similar across conditions (1.2 to 1.3) and the lowest stimulus frequency close to or above 0.85 kHz. For the resolved conditions, the same harmonic numbers could be used in both regimes. For the unresolved conditions, the low-frequency regime required the lowest possible pair of unresolved harmonics, *n* = 10 and *m* = 13: higher harmonic numbers would have required such low fundamental frequencies that the DPOAE templates would have fallen into a range in which speech-like DPOAEs cannot be measured reliably. The transitional regime allowed higher harmonic numbers, *n* = 15 and *m* = 18, which even produced slightly smaller correlations between stimulus waveforms and templates than the pair used in the low unresolved condition.

For each condition, the two waveforms with the lower harmonic number *n*, one from each voice, were summed to form *W* ^(1)(^*t*), and the two with the higher harmonic number *m* to form *W* ^(2)(^*t*). The two sums were delivered through separate loudspeakers to avoid distortion between the lower and the higher harmonic produced by the hardware.

### 2.5 Experimental setup

The experiment took place in a sound-proof, semi-anechoic chamber. Sounds were presented at a sampling rate of 44.1 kHz through a sound card (Fireface 802, RME) and an extended-bandwidth OAE probe system (ER10X, Etymotic Research), which provides one microphone and three loudspeakers per ear. Custom ear tips ensured a tight fit. Stimulus presentation and data acquisition were controlled by PsychoPy [33].

The audiobooks were presented to the right ear, and the waveforms *W* ^(1)(^*t*) and *W* ^(2)(^*t*) to the left ear, from which the speech-like DPOAEs analysed here were recorded. Unlike in JS25, *W* ^(1)(^*t*) and *W* ^(2)(^*t*) were also presented to the right ear, so that speech-like DPOAEs could additionally be recorded ipsilateral to the speech. In the right ear, the audiobook and the two stimulus waveforms were played through the three separate loudspeakers of the probe. The ipsilateral recordings require different analysis methods and will be reported separately; the present study is based exclusively on the recordings from the left ear, which are directly comparable to those of JS25.

The stimulus waveforms were presented to the left ear at an average sound pressure level of 57 dB SPL, measured over each two-minute trial. In the right ear, the combined level of the audiobooks and the stimulus waveforms was 69 dB SPL on average, which ensured that the stimulus waveforms were masked. Owing to an error in the calibration procedure that was discovered only after data collection, both levels exceeded the intended values of 50 and 60 dB SPL. The same error had caused the levels of JS25 to be reported too low: re-estimated with the corrected procedure, its stimulus and audiobook levels were both 57 dB SPL on average [25]. The stimulus level in the left ear was therefore the same in both studies, whereas the level in the right ear was about 12 dB higher here. All participants reported the levels as comfortable, none found either side too loud, and all reported that they could not hear the stimulus waveforms in the right ear. Possible effects of the levels on the middle-ear muscle reflex and the MOC reflex are considered in Section 4.5.

### 2.6 Procedure

The experiment began with a two-minute measurement of pure-tone DPOAEs, which confirmed that DPOAEs could be recorded in all participants. The main part consisted of four blocks, one per condition, in randomised order. Each block contained six trials of about two minutes, three with the female and three with the male voice as the target, in randomised order, so that each participant completed 24 trials. During each trial, participants looked at an image representing the attended story in the centre of the screen, to minimise eye movements. After each trial, they answered three comprehension questions about the attended story and rated their listening effort on a 13-point Likert scale. Instructions were presented on the screen, and responses were given by mouse click.

To verify that the measurement equipment itself did not produce speech-like DPOAEs, we obtained out-of-ear control recordings with an otherwise identical procedure, with the probe placed in the centre of the room, away from reflecting surfaces [25]. No speech-like DPOAEs emerged in these recordings.

### 2.7 Analysis of speech-like DPOAEs

All recordings were analysed with the Python pipeline of our methods paper, in which the metrics used here are described and compared in detail [25]. For each trial, the delay introduced by the equipment was estimated by cross-correlating the recording with the two stimulus waveforms, and then removed.

#### 2.7.1 Time domain: cross-correlation

For each trial and template *w*_2_*_n−m_*(*t*), we computed the complex cross-correlation between the recording and the analytic representation of the DPOAE template [16]. Its absolute value, the envelope *E*(*τ*), does not depend on the phase of the emission. The envelope was smoothed with a zero-phase low-pass filter and retained for lags *τ* between*−*0.5 and 0.5 s. Lags of 100 ms *≤ |τ| ≤* 500 ms, at which no emission is expected, served as the noise region, and the mean envelope within this region as the noise floor *µ_N_*.

Grand averages (GAs) were obtained by averaging the envelopes of all trials of all participants, separately for each stimulus type and attentional condition. The GA peak was located within *±*15 ms and described by a Gaussian fit with a constant baseline. The centre *τ^∗^* of the fit served as the latency of the emission and as the reference lag for the metrics below. As a measure of the strength of a GA peak, we report its signal-to-noise ratio (GA SNR), defined like the SNR of single trials below: 20 log_10_ of the GA peak value divided by the mean *µ_N_* of the GA in the noise region.

A GA peak was considered significant if it met two criteria [25]: it had to exceed the maximum of the GA in the noise region, and it had to lie at least four standard deviations of the GA in the noise region above the mean of this region. The first criterion alone is lenient for broad, smooth envelopes such as those of the present study, because neighbouring lags are then strongly correlated, so that the noise region contains only a few independent values. The second criterion does not depend on the number of independent values. To assess how consistently the emissions occurred across trials, we additionally considered the peak of a single trial (within *±*25 ms) significant if it exceeded the 97th percentile of the envelope in the noise region of that trial. This criterion is looser, because the envelope of a single trial is considerably noisier than a GA.

To compare the attentional conditions, we derived three metrics per participant from the cross-correlation. All of them were evaluated at the reference lag *τ^∗^* of the respective stimulus type and attentional condition, rather than at the optimal lag that yielded the maximal correlation in each trial, which could bias the results.

- *Peak magnitude.* The value of the envelope at *τ^∗^*, averaged across the trials of a participant. This is the amplitude analysed in JS25. Because the envelope is the magnitude of a complex number, the peak magnitude contains a positive contribution from noise even when no emission is present.
- *Signal-to-noise ratio (SNR).* The ratio 20 log10 [E(*τ^∗^*)/µN]per trial, averaged across the trials of a participant. The SNR relates each trial to its own noise floor and was the most consistent of the cross-correlation metrics in our methods paper [25]; we therefore use it as the main time-domain metric. Note, however, that the noise floor also contains chance correlations between the DPOAE template and the other waveforms of the recording, including the stimuli of the competing voice.
- *Area under the curve (AUC).* With the narrowband templates of the present study, the GA peaks were an order of magnitude broader than in JS25 (Section 3.2). The energy of the emission is then distributed over many lags, and the value at a single lag captures only part of it. We therefore also computed the area of each participant’s GA envelope, averaged across the trials of this participant, above its noise floor within *±σ*_ac_ around *τ^∗^*, where *σ*_ac_ is the width of the auto-correlation of the DPOAE template. Values below the noise floor were treated as missing.

#### 2.7.2 Frequency domain: magnitude-squared coherence

The narrow bandwidth of the DPOAE templates has opposite consequences in the time and in the frequency domain. By the time–frequency uncertainty principle, a narrowband template has a long auto-correlation, which causes the broad cross-correlation peaks and the poor temporal resolution of the present study (Section 3.2). In the frequency domain, in contrast, the same templates concentrate their power in narrow bands of a few tens of hertz around their mean frequencies (Table 2), which can be resolved well with long analysis segments. A frequency-domain measure may therefore capture more information about the emission than the time-domain metrics, and it does not require a reference lag. We therefore computed the magnitude-squared coherence (MSC) between template and recording, using Welch’s method with Hann-windowed segments of 16,384 samples (371.5 ms) and 50 % overlap. The MSC states which fraction of the power of the recording at a given frequency can be explained by a linear transformation of the DPOAE template. We corrected it for the positive bias that arises from estimating it from a finite number of segments and averaged it across the frequency band of the DPOAE template, defined as the band containing the central 90 % of the DPOAE template’s power. In this average, each frequency was weighted by the power of the DPOAE template, so that frequencies at which the DPOAE template, and therefore presumably the emission, carries more power contribute more strongly. The resulting power-weighted MSC was computed for every trial and averaged across the trials of a participant.

#### 2.7.3 Attentional modulation coefficient

To remove factors that scale the attended and the ignored state of a participant equally, such as the overall strength of the emissions or the fit of the probe, we also computed the attentional modulation coefficient (AMC) [16],

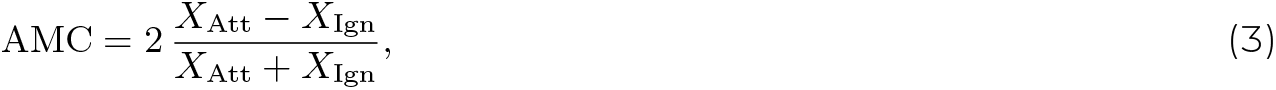

where *X*_Att_ and *X*_Ign_ are the values of a DPOAE metric obtained when the participant attended or ignored the corresponding voice, respectively. Positive values indicate larger values when the voice was attended. We computed the AMC for the peak magnitude, the AUC and the power-weighted MSC, but not for the SNR: the SNR is already referenced to the noise floor, and since it can be close to zero or negative on the decibel scale, Eq. (3) would be unstable.

### 2.8 Statistical analysis

Attended and ignored states were compared separately for each stimulus type. Values more than 1.5 interquartile ranges below the first or above the third quartile were excluded as outliers; for paired comparisons, the exclusion was applied pairwise, so that participants remained matched. Paired *t*-tests were used if both samples passed a Shapiro– Wilk test of normality, and Wilcoxon signed-rank tests otherwise. For the AUC, values below the noise floor left too few complete pairs for some stimulus types, so that the two states were compared with unpaired *t*-tests or Mann–Whitney *U* tests. AMCs were tested against zero with one-sample *t*-tests or Wilcoxon signed-rank tests, chosen in the same way. For each metric, *p*-values were corrected across stimulus types with the Benjamini–Hochberg procedure [34]: across the seven stimulus types with a significant GA peak (Section 3.2) for the metrics derived from the cross-correlation, and across all eight stimulus types for the MSC. Comprehension scores and listening effort were averaged per participant, condition and target voice; the two target voices were compared within each condition with the same paired tests, corrected across the four conditions.

#### Sensitivity of the design

To assess whether the present design could have detected an effect of the size observed in JS25, we used the single-trial amplitudes of the stimulus F_res_ of JS25 in the conditions Att. F and Att. M, with the same outlier exclusion as above (*N* = 36). Since the present study used three instead of six trials per condition, we estimated the effect size that JS25 would have yielded with three trials by subsampling. For every participant, three of the six trials per condition were drawn at random, and the paired effect size *d_z_* (mean difference divided by the standard deviation of the differences) was computed; this was repeated 2,000 times. With the median of these effect sizes, we computed the power of a two-sided paired *t*-test with *N* = 44. We report the power for statistical significance at *α* = 0.05 and at *α* = 0.05*/*7, which bound the power of the Benjamini–Hochberg procedure: its threshold for a single effect is *α/*7 if only one of the seven stimulus types tested with the cross-correlation metrics carries an effect, and approaches *α* as more stimulus types carry one. Finally, we used equivalence tests to determine for each stimulus type and metric whether an effect of size *|d_z_|* could be excluded: that is the case when the 90% confidence interval of the observed effect size lies entirely within *±d_z_*. This is equivalent to rejecting both one-sided null hypotheses that the observed effect size is both smaller than *−d_z_* and bigger than +*d_z_* in two one-sided tests (TOST, *p_TOST_ <* 0.05) [35].

### 2.9 Post hoc model of the cochlear excitation by the two voices

A frequency-specific modulation of the cochlear amplifier through the MOC system can only favour an attended voice if its harmonics excite other cochlear places than those of the competing voice. Since the pitch shifting reduced ΔERB from 2.6 in JS25 to about 1 here (Table 1), we estimated, for both studies, at which places the two voices excite the cochlea in the same way. This analysis was conducted post hoc, after the effect of JS25 had not been reproduced, to explore whether the closer spacing of the voices could account for the absence of the effect.

Each voice was represented by a steady harmonic complex at its mean fundamental frequency (Table 1; 195 and 90 Hz for the female and the male voice of JS25), with all harmonics up to 5 kHz. The power of the harmonics fell by 6 dB per octave, a common idealisation of the spectrum of voiced speech [36]. The complex was filtered with symmetric rounded-exponential auditory filters [24], so that the excitation at the cochlear place with characteristic frequency *f_c_* was

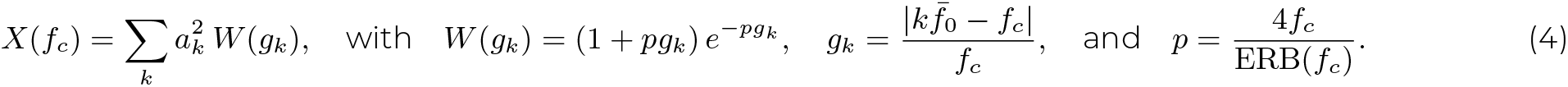

Hereby *a*^2^_*k*_ is the power of the *k*-th harmonic, *g_k_* the distance of its frequency *kf̄*^0^ from the characteristic frequency, relative to *f_c_*, *W* (*g_k_*) the weight of the filter at the relative distance *g_k_*, *p* the parameter that sets the sharpness of the filter, and ERB(*f_c_*) = 24.7 (4.37 *f_c_/*1000 + 1) is the equivalent rectangular bandwidth of the filter, that is, its width in Hz for *f_c_* in Hz [24]. This bandwidth should not be confused with the ERB number *E*(*f*) in Eq. (1), which counts how many such bandwidths fit below the frequency *f*. It is obtained by integrating 1*/*ERB(*f*), which gives Eq. (1) its logarithmic form. The excitation was evaluated at places spaced evenly on the ERB-number scale between 0.1 and 4.5 kHz, expressed in dB and normalised to the larger maximum of the two voices. As long as the harmonics of a voice are resolved, its excitation pattern shows a peak at the place of each harmonic; where several harmonics fall into one filter, the pattern becomes flat.

We defined shared places as places at which the peaks of both voices coincide. Before the patterns were compared, a moving average over 3 ERB was subtracted from each of them. This removes slow changes of the excitation level along the cochlea, which mainly stem from the spectral slope of the voice, and leaves the ripple produced by the individual harmonic peaks. Without this step, two patterns that both fall towards the base would correlate even where their peaks alternate. A place was considered shared if two conditions were met: the local correlation between the two detrended patterns within a window of 2 ERB exceeded 0.5, and both patterns showed a harmonic ripple of more than 0.5 dB (local root-mean-square of the detrended pattern within the same window). The second condition excludes places at which flat, unresolved patterns correlate by chance. With this ripple threshold, a voice with a fundamental frequency between 100 and 200 Hz counts as resolved up to about its seventh to eighth harmonic, and a voice with a lower fundamental frequency up to about its sixth to seventh harmonic. This is slightly more conservative than the common limit of about the tenth harmonic, so that only clearly resolved peaks count as shared. As a summary measure, we computed the percentage of shared places between 0.5 and 4 kHz.

The model is deliberately simple. It is linear and does not depend on level, so that it includes neither cochlear compression and suppression nor any action of the efferent system. It neglects the fluctuations of the fundamental frequency over time as well as the formant structure of real speech, and it assumes equal levels for both voices and a fixed spectral slope; the shared regions were, however, essentially unchanged for slopes between 0 and *−*12 dB per octave. The criteria that define a shared place are, to some degree, arbitrary.

## 3 RESULTS

### 3.1 Behavioural data

Participants answered most comprehension questions correctly, with mean scores between 89 and 94 % across conditions and target voices, and comprehension did not differ between the two target voices in any condition (all corrected *p >* 0.89). Listening effort, however, was rated higher by 1.3 to 1.9 points on the 13-point scale when the male voice was attended, in all four conditions (all corrected *p <* 10*^−^*^5)^. This matches JS25, in which attending the male voice was also rated as more effortful. Listening effort was highest in the transitional resolved condition, for both target voices (female: 7.7 *±* 1.8; male: 9.6 *±* 1.6; mean *±* SD). In this condition, the fundamental frequencies were shifted furthest, to 270 and 325 Hz, close to the highest values at which the speech remained intelligible during stimulus development, and participants described the two voices as harder to tell apart. Since comprehension remained high and did not differ between the voices, we conclude that participants were able to follow the instructed voice in all conditions.

### 3.2 Speech-like DPOAEs

The GAs of the cross-correlation envelopes showed clear peaks near zero lag for most stimulus types (Fig. 2B), and 14 of the 16 GAs met both significance criteria. The strongest peaks occurred for TM_unres_, with GA SNRs of 17.4 and 17.6 dB in the attended and the ignored state (87 and 103 standard deviations above the noise floor), and for TF_unres_ (10.9 and 10.0 dB; 32 and 21 standard deviations). The weakest significant peaks were those of the male voice in the low-frequency regime: the GAs of LM_res_ (2.2 and 3.1 dB) and of LM_unres_ in the attended state (2.5 dB) lay only 4.7 to 5.0 standard deviations above the noise floor. All other significant peaks, including that of LM_unres_ in the ignored state (4.4 dB; 8.7 standard deviations), lay at least 8.4 standard deviations above it. For LF_unres_, the GA showed no clear peak in either state (2.5 and 1.5 dB; 2.7 and 1.3 standard deviations) and did not meet the significance criteria; in the ignored state, it did not even exceed the maximum of the noise region. Since all metrics derived from the cross-correlation are evaluated at the GA peak, LF_unres_ was excluded from their analysis.

**Figure 2:**
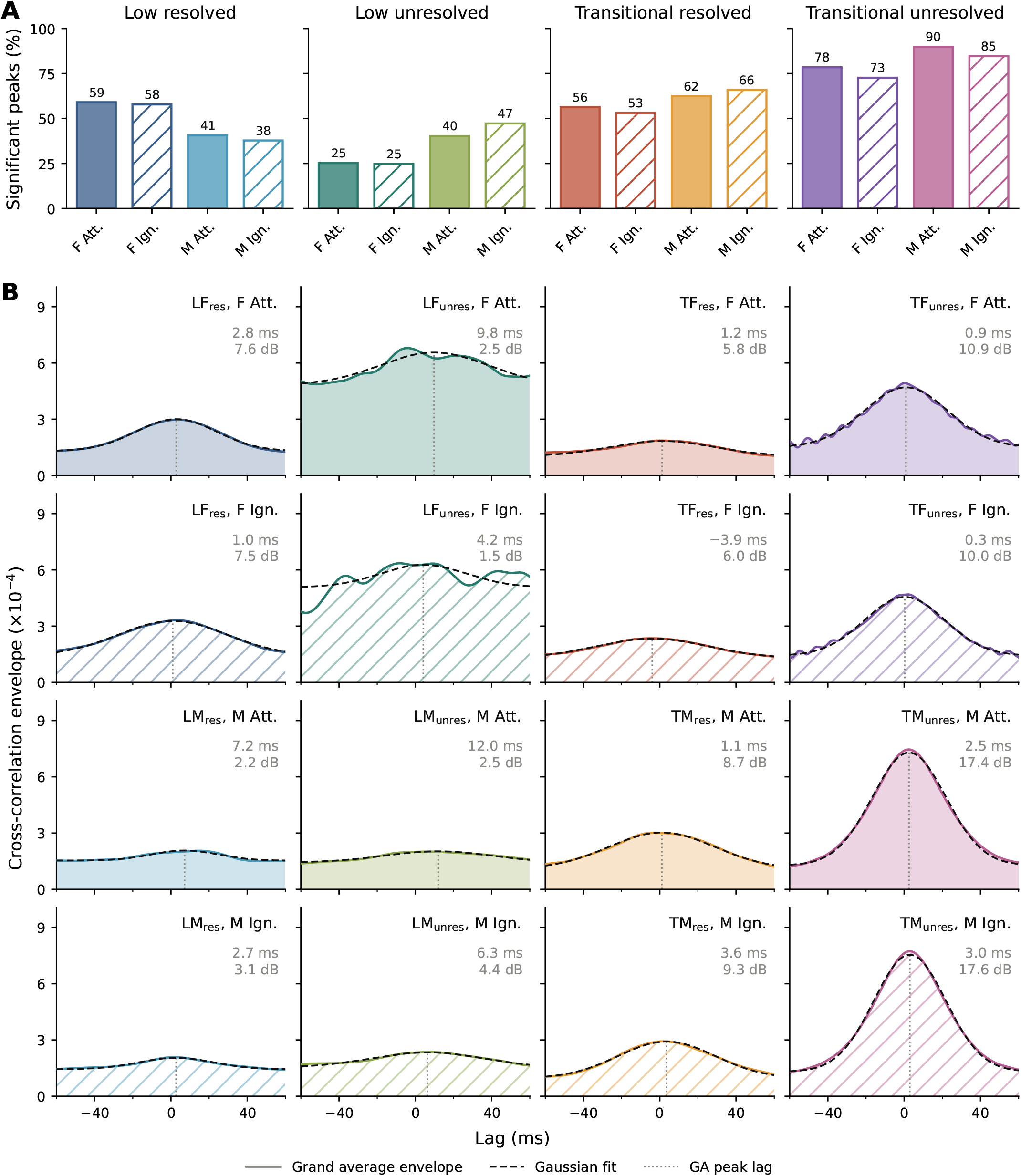
Speech-like DPOAEs in the time domain. (A) Percentage of trials with a significant peak, for the four conditions and the female (F) and male (M) voice in the attended (Att., filled) and ignored (Ign., hatched) state. (B) Grand average envelopes of the complex cross-correlation (coloured lines) with fitted Gaussians (dashed lines). The dotted vertical lines mark the fitted peak lag; the grey numbers give the peak lag and the GA SNR (GA peak relative to the mean of the GA at lags of 100 to 500 ms). L, T: low and transitional frequency regime; F, M: female and male voice; res, unres: resolved and unresolved harmonics.

All peaks were broad, with fitted widths *σ* between 17 and 30 ms, compared with 1.5 to 3.9 ms in JS25 [25]. As expected for narrowband templates, the widths closely matched those of the auto-correlations of the DPOAE templates. The fitted peak lags ranged from -3.9 to 12.0 ms; given the widths of the peaks, we regard the differences in latency between the stimulus types as random fluctuations rather than systematic trends.

The proportion of trials with a significant peak followed the same pattern (Fig. 2A): it was highest for the transitional unresolved stimuli (73–90 %), intermediate for the resolved stimuli (38–66 %), and lowest for the low unresolved stimuli (25–47 %). It did not differ systematically between the attentional conditions.

### 3.3 No consistent effect of attention

#### Peak magnitude

We first analysed the peak magnitude, the metric in which JS25 had shown the attentional effect. It did not differ between the attended and the ignored state for any of the seven tested stimulus types (all corrected *p ≥* 0.39), and its AMC did not differ from zero either (all corrected *p ≥* 0.08). The peak magnitude and its AMC are therefore not shown in the figures.

#### SNR

The SNR differed between the attentional states for two stimulus types, in opposite directions (Fig. 3A). For LM_unres_, it was higher when the male voice was ignored than when it was attended (median 4.7 vs. 1.7 dB; corrected *p* = 0.002). For TF_unres_, it was higher when the female voice was attended (median 9.6 vs. 8.4 dB; corrected *p* = 0.017). No other stimulus type showed a difference (all corrected *p ≥* 0.77).

**Figure 3:**
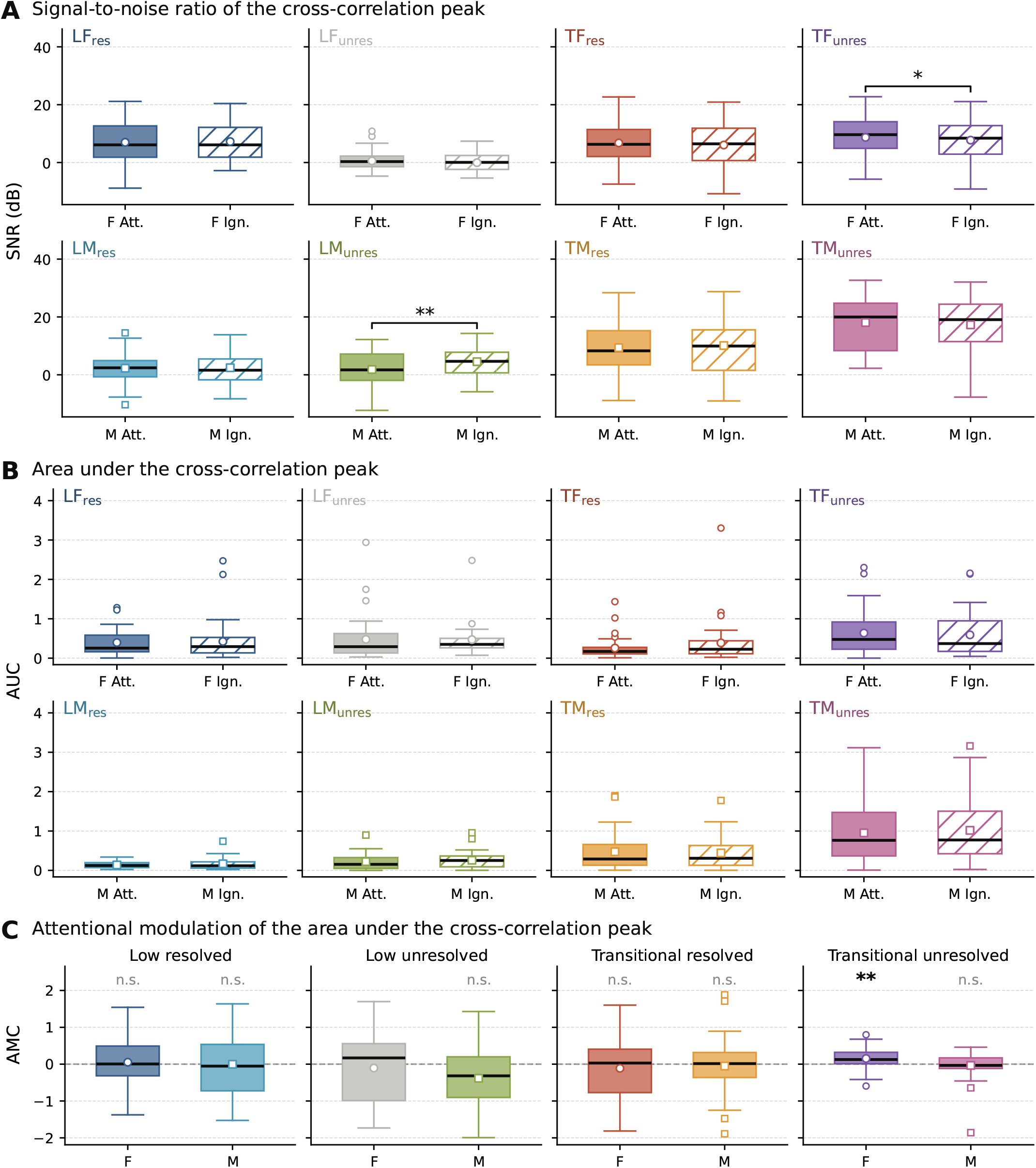
Time-domain metrics in the attended (Att., filled) and the ignored (Ign., hatched) state, with one value per participant. (A) SNR of the cross-correlation at the GA peak lag. (B) Area under the cross-correlation peak above the noise floor. (C) Attentional modulation coefficient (AMC) of the AUC; positive values indicate larger values when the corresponding voice was attended. Circles and squares mark the means for the female and the male voice. LF_unres_ is shown in grey and was not tested, because its GA showed no significant peak (Section 3.2). Significance after Benjamini–Hochberg correction across the seven tested stimulus types: *^∗p^ <* 0.05, *^∗∗p^ <* 0.01; n.s.: not significant.

#### AUC

The AUC itself did not differ between the attentional states for any stimulus type (Fig. 3B; all corrected *p ≥* 0.92). Its AMC, which relates the two states of each participant to each other, was positive for TF_unres_ (median 0.13; corrected *p* = 0.002; Fig. 3C), in the same direction as the SNR. For LM_unres_, the AMC was negative, again in the same direction as the SNR, but did not survive the correction for multiple comparisons (median *−*0.32; uncorrected *p* = 0.033, corrected *p* = 0.11). The AMCs of all other stimulus types were close to zero (all corrected *p ≥* 0.68).

#### Frequency domain

Since the MSC does not require a GA peak, there was no formal reason to exclude LF_unres_ from the frequency-domain analysis, and we included it for completeness. Its results should nevertheless be read with caution: its GA showed no significant peak (Section 3.2), and its MSC did not separate in-ear from out-of-ear recordings in our methods paper [25], so that a reliable emission cannot be expected for this stimulus type. After correction for multiple comparisons, the power-weighted MSC did not differ between the attentional states for any stimulus type (Fig. 4A; all corrected *p ≥* 0.07). The smallest *p*-value occurred for LM_unres_, with larger values in the ignored state, in the same direction as the SNR (uncorrected *p* = 0.010, corrected *p* = 0.077). The AMC of the power-weighted MSC did not differ from zero for any stimulus type after correction either (Fig. 4B). It came closest to significance for the two low-frequency unresolved stimuli, with opposite signs: it was negative for LM_unres_ (median *−*0.45; uncorrected *p* = 0.007, corrected *p* = 0.052) and positive for LF_unres_ (median 0.65; uncorrected *p* = 0.014, corrected *p* = 0.058), although, for the reasons given above, the latter is unlikely to reflect the modulation of an emission.

**Figure 4:**
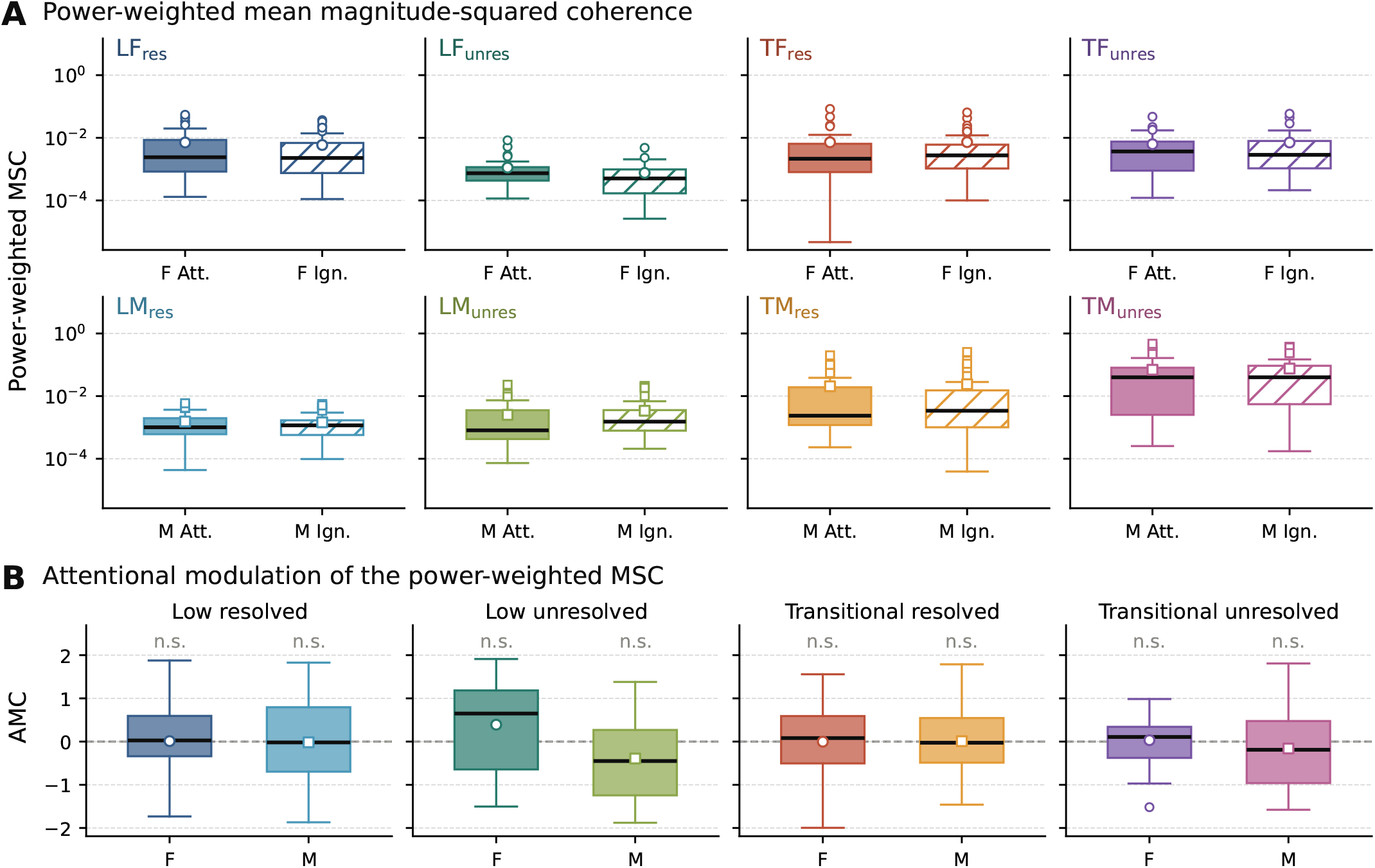
Frequency-domain metrics. (A) Power-weighted, bias-corrected mean magnitude-squared coherence (MSC) between template and recording in the attended (Att., filled) and the ignored (Ign., hatched) state, with one value per participant; note the logarithmic axis. (B) AMC of the power-weighted MSC. None of the comparisons was significant after Benjamini–Hochberg correction across the eight stimulus types.

Both the AUC and the power-weighted MSC showed several high outliers (Figs. 3B and 4A). These reflected participants with particularly strong emissions rather than noise, and they were largely consistent across metrics: 26 of the 28 outliers of the AUC and 72 of the 77 outliers of the MSC belonged to the quarter of participants with the highest SNR for the respective stimulus type and attentional state, and 20 of the outliers of the AUC were also outliers of the MSC.

Since the AUC and the MSC are expressed on a linear scale, whereas the SNR is expressed in decibels, a few strong participants stand out more clearly in them.

In summary, no metric showed an effect of attention that was consistent across stimulus types, voices or metrics. The significant effects were restricted to two stimulus types: LM_unres_, with larger values when the male voice was ignored, and TF_unres_, with larger values when the female voice was attended. They appeared in some metrics but not in others, and none of them appeared in the peak magnitude, the metric in which the effect of JS25 had been found.

### 3.4 Sensitivity of the design

In JS25, the effect for F_res_ amounted to *d_z_* = 0.82 when computed with all six trials per condition. In this study, we utilized only three trials for each condition. With three randomly selected trials per condition out of the six from JS25, the median effect size decreased to *d_z_* = 0.53 (90 % of the subsamples between 0.32 and 0.72). Halving the number of trials thus reduced the effect size by a factor of about 1.5, which shows that the trial-to-trial variability in JS25 was large relative to the variability between participants. The reduction even slightly exceeds the factor of *^√^*^2^ expected if all variability stemmed from single trials, presumably because single trials with extreme values weigh more heavily in a mean of three trials.

An effect of *d_z_* = 0.53 would have been detected with a power of 0.75 at the strictest threshold of the Benjamini–Hochberg procedure (*α* = 0.05*/*7) and with a power of up to 0.93 if several stimulus types had carried an effect (*α* = 0.05). The power for a single affected stimulus type thus lay slightly below the conventional value of 0.8, mainly because only three trials per condition were recorded. Whether an effect of the size observed in JS25 can be excluded was therefore assessed directly with equivalence tests.

While the comparisons above tested for evidence of an effect of attention, the equivalence tests ask whether an effect as large as in JS25, *|d_z_| ≥* 0.53, can be excluded. This is particularly informative where no significant effect was found, since the absence of a significant difference alone does not show that an effect is absent. An effect of size *|d_z_| ≥* 0.53 can be excluded when the 90 % confidence interval of the observed effect size lies entirely within *±*0.53 (two one-sided tests, TOST, *p*_TOST_ *<* 0.05). Together, the two kinds of test sort the 29 combinations of stimulus type and metric (seven stimulus types for each of the three cross-correlation metrics and eight for the MSC) into three groups.

First, for 22 combinations, an effect as large as in JS25 could be excluded. In these combinations, any effect of attention was either considerably smaller than in JS25 or absent; the data cannot distinguish between the two. They included six of the seven stimulus types for the peak magnitude, the metric of the JS25 effect, and 15 of the 16 combinations of the two resolved conditions, which correspond most closely to the stimulus F_res_ of JS25.

Second, for three combinations, an effect of this size could not be excluded, and an effect of attention was found: the SNR of LM_unres_ (*d_z_* = *−*0.61) and of TF_unres_ (*d_z_* = 0.45), and the AUC of TF_unres_ (*d_z_* = 0.55). For the AUC, the effect was significant only in its AMC, which relates the two states of each participant to each other; the unpaired comparison of the AUC values themselves was not significant.

Third, for the remaining four combinations, the result was inconclusive: an effect of the size observed in JS25 could neither be excluded nor demonstrated. For LM_unres_, the effect sizes of the peak magnitude, the AUC and the MSC were negative (*d_z_* between *−*0.32 and *−*0.39), in the same direction as its SNR effect, but too small to be significant and too uncertain to be excluded. For LF_res_, the AUC was based on fewer participants, because values below the noise floor were missing, and its confidence interval was correspondingly wide.

Taken together, the present study was sensitive enough to detect an effect of the size observed in JS25.

### 3.5 Excitation patterns of the two voices

Figure 5 shows the excitation patterns of the two voices obtained with the post hoc model (Section 2.9), for JS25 and for the four conditions of the present study.

**Figure 5:**
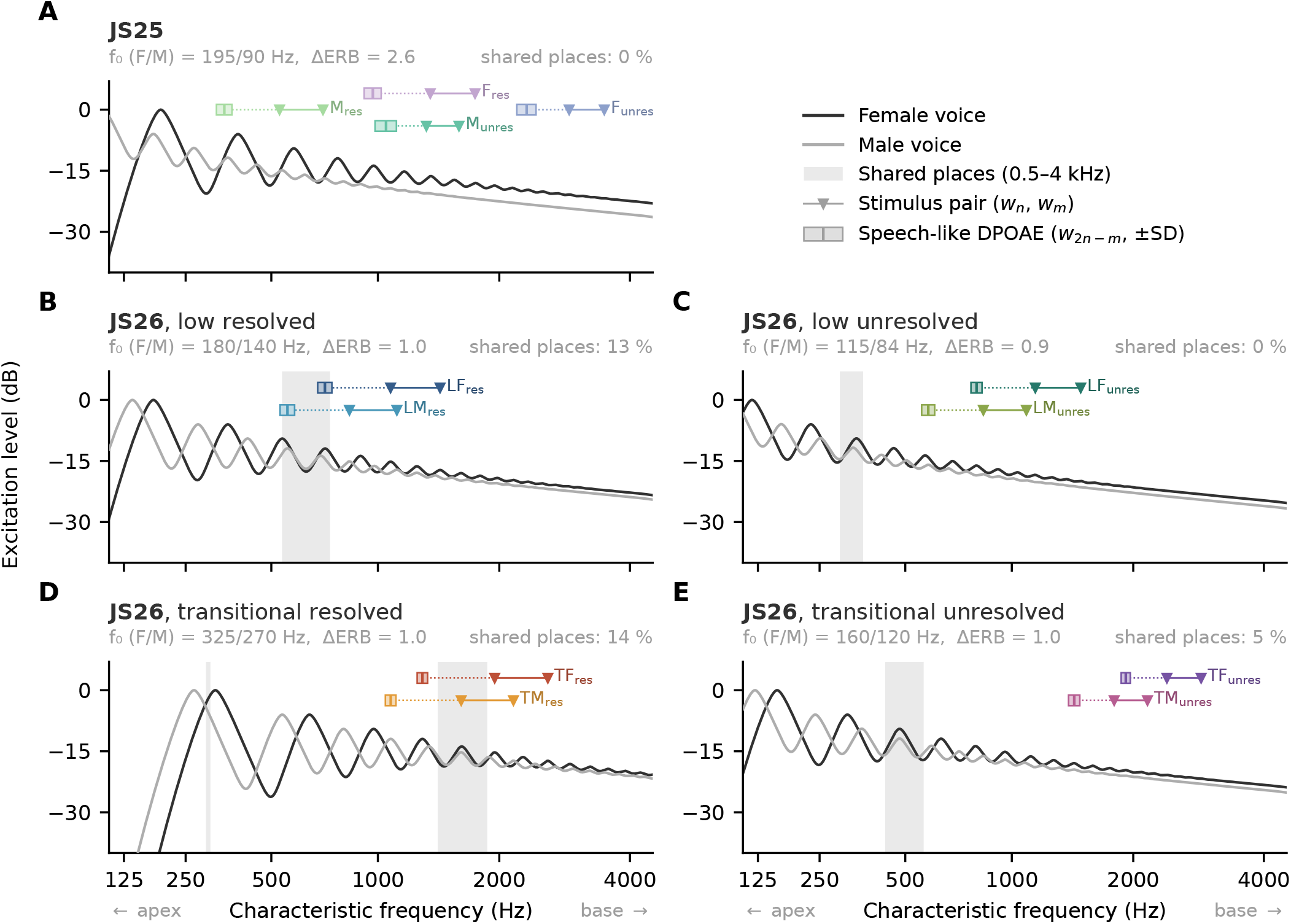
Excitation patterns of the female (black) and the male (grey) voice along the cochlea, for JS25 (A) and for the four conditions of the present study (B–E), obtained with the post hoc model (Section 2.9). Grey bands mark shared places, at which the harmonic peaks of both voices coincide; the percentage of shared places between 0.5 and 4 kHz is given above each panel. Only this range was counted, because all stimulus and emission frequencies lie within it; shared places at lower frequencies, such as those in the low unresolved condition (C), are shown for completeness but do not enter the percentage. Coloured markers show the mean frequencies of the stimulus pairs (*w_n_*, *w_m_*; triangles) and of the corresponding speech-like DPOAEs (*w*_2_*_n−m_*; squares, with *±*SD; cf. Table 2).

In JS25 (Fig. 5A), the male voice, with its low fundamental frequency of 90 Hz, showed pronounced harmonic peaks only up to about 0.6 kHz, and its excitation pattern became essentially flat above about 1 kHz. The female voice, in contrast, produced distinct peaks up to about 2 kHz. There were no shared places between 0.5 and 4 kHz. The stimuli and the emission of F_res_ therefore lay at places at which only the female voice showed harmonic peaks. The same held for the stimulus M_unres_, whose mean stimulus and emission frequencies almost coincided with those of Fres.

In the present study, both voices produced resolved peaks of similar spacing, which alternated along the cochlea and coincided in some regions (Fig. 5B–E). In the two resolved conditions, which had the most shared places (13 % in the low and 14 % in the transitional resolved condition, between 0.5 and 4 kHz), these shared regions lay at or close to the stimuli and the emissions. In the low resolved condition, the shared region spanned about 0.54 to 0.74 kHz (Fig. 5B) and contained the emission frequencies of both voices, at about 0.72 kHz for LF_res_ and 0.56 kHz for LM_res_. In the transitional resolved condition, it spanned about 1.4 to 1.9 kHz (Fig. 5D); it contained the lower stimulus waveform of TM_res_ and lay just above the emission of TF_res_ at 1.3 kHz. In the two unresolved conditions, the shared regions lay well below the stimuli and the emissions, at about 0.30 to 0.36 kHz in the low unresolved and 0.43 to 0.57 kHz in the transitional unresolved condition (Fig. 5C, E), so that only 0 and 5 % of the places between 0.5 and 4 kHz were shared. Thus, LM_res_ and LM_unres_, whose stimulus and emission frequencies were almost identical (Table 2), differed in one respect: the emission of LM_res_ lay in a shared region, that of LM_unres_ did not.

## 4 DISCUSSION

We set out to reproduce the attentional modulation of speech-like DPOAEs reported in JS25 [17], in a design that separated frequency region and harmonic resolvability. Participants followed the instructed voice, and speech-like DPOAEs emerged for most stimulus types, although those of the male voice in the low-frequency regime were close to the detection limit and LF_unres_ yielded no measurable emission. Attention, however, did not significantly modulate the amplitude of the emissions for any stimulus type. Further metrics revealed isolated effects for two stimulus types, LMunres and TF_unres_, which went in opposite directions and were not consistent across metrics. The power analysis and the equivalence tests show that this outcome cannot be explained by a lack of sensitivity alone: an effect of the size observed in JS25 would most likely have been detected, and it can be ruled out for most combinations of stimulus type and metric.

The present findings thus contrast with those of JS25 and, to a lesser degree, with those of Saiz-Alía et al. [16]. We do not consider either study to invalidate the other. The effect in JS25 was strong and unambiguous for one stimulus type, and we see no flaw in its design that could have produced it artificially. At the same time, the changes made for the present study were intended as refinements, and it was not obvious at first why they should have abolished a genuine effect. In the following, we argue that one of these changes, the pitch shifting and the resulting much smaller separation of the two voices, may have done exactly that, and that the narrower bandwidth of the stimuli may have amplified its consequences. A further difference from JS25, the higher level of the speech in the ear contralateral to the recordings, is considered in Section 4.5.

### 4.1 Overlap of the two voices along the cochlea

Separating the stimuli of the two voices in frequency, as we did in the present design, does not separate the voices themselves. Each voice excites the cochlea with all of its harmonics, not only with the two from which a stimulus pair is derived, and a frequency-specific modulation through the MOC system can only favour the attended voice at places that this voice does not share with the competing one. The model suggests that the pitch shifting changed exactly this (Section 3.5). In JS25, the two voices were separated by 2.6 ERB, and the harmonics of the low male voice were unresolved above about 1 kHz. The resolved harmonics of the female voice therefore had large parts of the cochlea to themselves, including the places of the stimulus F_res_, which showed the strong effect. With a separation of only about one ERB, in contrast, both voices produced resolved harmonics of similar spacing, which coincided in some regions along the cochlea. Whether an emission can reflect a modulation directed at one voice then depends on where its stimuli and its emission lie relative to these shared regions, and the results of the present study follow this distinction closely.

In the low resolved condition, which was designed to reproduce the effect of JS25, the emissions of both voices lay in a shared region. Attention to either voice would then modulate both emissions in the same way, and since the attended state of one voice was always recorded together with the ignored state of the other, the difference between the two states would vanish for both voices. In the transitional resolved condition, the shared region contained a stimulus waveform of one voice and lay close to the emission of the other. Since distortion products presumably arise where the excitation patterns of the two stimulus waveforms overlap and are additionally reflected at their own characteristic place [37], both the stimulus and the emission frequencies may matter here. Neither resolved condition showed an attentional effect. The isolated effects, in contrast, emerged in the two unresolved conditions, in which the shared regions lay well away from the stimuli and the emissions.

The contrast between LM_res_ and LM_unres_ points in the same direction. Both probed almost the same region of the cochlea, but only for LM_unres_ was this region not shared with the female voice, and only LM_unres_ showed an effect. Its emission became weaker when the male voice was attended, which is the direction of the effect in JS25. This comparison is, however, limited by the weak emissions of both stimulus types (Section 3.2). The model is also consistent with our interpretation of the effect for Munres in JS25, which we had attributed to its overlap with F_res_ [17]: its stimuli and emission lay in the region in which only the female voice showed harmonic peaks, so that a modulation directed at the female voice would have affected M_unres_ as well.

Not all observations fit this account equally well. For LF_unres_, the measurement itself failed (Section 3.2), so that the absence of an effect is not informative. TM_unres_, whose stimuli and emission did not lie in a shared region, showed no effect, although it produced the strongest emission of the study. At these higher frequencies, attentional effects may be weaker or less consistent; in JS25, the stimulus F_unres_ at similar frequencies did not show an effect either (Section 4.3). Moreover, the account is post hoc, and the model is deliberately simple (Section 2.9). In particular, it neglects that *f*_0_ fluctuates over time, which continuously shifts the shared places. In addition, the frequency selectivity of the MOC system in humans is not well characterised [38–40].

Nevertheless, a single feature of the stimulus design could explain both the absence of the effect in the condition that was designed to reproduce JS25 and the location of the isolated effects. The account also makes a testable prediction: the effect of JS25 should re-emerge when the two voices are separated sufficiently for the stimuli and emissions of both voices to lie outside shared places.

### 4.2 Bandwidth of the stimuli

The consequences of the overlap between the different stimuli in terms of cochlear excitation described above may have been amplified by the narrow bandwidth of the stimuli. Because the voices were close, we narrowed the filter that extracts the fundamental waveform, to reduce the correlation between the stimuli of one voice and the DPOAE templates of the other. As a result, the stimulus waveforms varied much less than the harmonics of the voices from which they were derived. The fluctuations of a harmonic scale with its harmonic number: for LM_res_, for instance, the sixth harmonic of the voice fluctuated with a standard deviation of about 170 Hz (6 *×* 28 Hz; Table 1), but the stimulus waveform *w*_6_(*t*) only with 42 Hz (Table 2). A stimulus pair whose generation sites happen to lie in a shared region therefore stays within this region most of the time. With the broader stimuli of JS25, the waveforms followed a larger part of the natural pitch contour and covered a wider range of cochlear places, so that the emission would have been less dominated by any single region. Broader stimuli might thus have partly compensated for an influence of the competing voice. In addition, narrowband stimuli follow the attended harmonics less closely: when the fundamental frequency of the voice moves away from its mean, the stimuli lag behind and probe places at which the attended voice might momentarily not be present.

The narrow bandwidth also had a methodological consequence. The DPOAE templates had long auto-correlations, so that the cross-correlation peaks were an order of magnitude broader than in JS25. A modulation that changes the shape rather than the height of the peak may then be missed by the peak magnitude, which was one reason to include the AUC and the MSC. These metrics, however, did not reveal a consistent effect either. Since the bandwidth is a property of the stimuli rather than of the analysis, its influence cannot be assessed retrospectively with the present or the previous data.

### 4.3 Direction of the isolated effects

The isolated effects went in opposite directions. For LM_unres_, the emission was larger when the corresponding voice was ignored, as for F_res_ in JS25; for TF_unres_, it was larger when the voice was attended, as reported by Saiz-Alía et al. [16]. In JS25, we had hypothesised that the direction of the effect depends on the frequency regime, with smaller emissions for attended voices only in the low-frequency, apical regime [17]. The two effects of the present study, one in each regime, would be consistent with this hypothesis. A purely inhibitory action of the MOC system, in contrast, would predict smaller emissions for the attended voice in both regimes, which the effect for TF_unres_ does not support.

Two points complicate this interpretation. First, both effects occurred for stimuli derived from unresolved harmonics, whereas JS25 showed an effect for resolved harmonics. For the low unresolved condition, however, the distinction between resolved and unresolved harmonics may be less clear-cut. To place unresolved harmonics into the low-frequency regime, we had to use the lowest possible harmonic numbers, *n* = 10 and *m* = 13, close to the transition between resolved and unresolved harmonics. At these harmonic numbers, adjacent harmonics were spaced by 0.73 to 0.77 auditory filter bandwidths at the frequency of the lower stimulus waveform (*f̄*^0^*/*ERB(*nf̄*^0^)), compared with 0.53 to 0.58 for the transitional unresolved stimuli and the unresolved stimuli of JS25. Moreover, the two waveforms of a pair were three harmonics apart, with a frequency ratio of 1.3, and were therefore resolved relative to each other in all conditions. Stimuli derived from neighbouring unresolved harmonics would avoid this, but in pilot recordings during stimulus development they did not yield measurable speech-like DPOAEs. The stimuli of LM_unres_ may thus have behaved more like resolved than like fully unresolved harmonics.

Second, the stimulus F_unres_ of JS25, with stimulus frequencies of about 2.9 and 3.5 kHz and an emission at about 2.3 kHz, was close to TF_unres_ of the present study (2.4 and 2.9 kHz, emission at 1.9 kHz), but did not show an attentional effect. Given these inconsistencies and the number of comparisons, we regard the direction of the isolated effects as a hint for future studies rather than as a finding.

### 4.4 Measurement quality in the low-frequency regime

The account above explains the absence of an attentional effect in the resolved conditions by the overlap of the voices. For LM_res_, a second reason may contribute. Together with that of LM_unres_ in the attended state, its GA peaks were the weakest significant peaks of the study, only 4.7 to 5.0 standard deviations above the noise floor, and only about 40 % of its trials showed a significant peak (Fig. 2). Since the metrics derived from the cross-correlation are evaluated at the GA peak, they carry a considerable uncertainty for these stimulus types, and an attentional modulation may simply have been masked by the variability of the measurement. The question raised by JS25, whether the resolved harmonics of the male voice are modulated by attention as well, therefore remains unanswered.

The same caution applies to the effect for LM_unres_. With templates at about 560 to 600 Hz, the emissions of the male voice in the low-frequency regime lay at the lower end of the frequency range in which speech-like DPOAEs could be measured here; in JS25, the stimulus M_res_, with a template at about 350 Hz, failed entirely [17]. For designs with several simultaneous stimulus pairs and narrowband templates, emission frequencies below about 600 Hz should therefore be avoided.

The failure of LF_unres_ does not fit into this picture, since its template lay at a higher frequency, about 805 Hz, close to the DPOAE template frequencies for which Saiz-Alía et al. [16] obtained clear speech-like DPOAEs. We did not expect this outcome. A possible explanation is the proximity of its template to a stimulus waveform of the competing voice: in the low unresolved condition, the lower stimulus waveform of the male voice lay at 840 *±* 26 Hz, only about 35 Hz above the DPOAE template of LF_unres_ at 805 *±* 28 Hz (Table 2), and was presented at a much higher level than the emission. Chance correlations between the DPOAE template and this stimulus waveform would raise the noise floor of the cross-correlation. Indeed, the noise floor of the GAs of LF_unres_ was 3.2 to 5.6 times higher than that of any other stimulus type (5.1 and 5.3 *×* 10*^−^*^4,^ compared with 0.96 to 1.6 *×* 10*^−^*^4)^. In addition, a strong stimulus close to the emission frequency may suppress the emission itself. In the study of Saiz-Alía et al. [16], only one stimulus pair was presented at a time, so that no such neighbouring waveform was present. Whatever the cause, this case shows that, with several simultaneous stimulus pairs, the DPOAE templates should be kept well away not only from the waveforms of their own stimulus pair but also from those of the competing voice.

### 4.5 Possible influence of the middle-ear muscle and MOC reflexes

Owing to the calibration error described in Section 2.5, the sound levels were higher than intended in both ears. This raises the question whether the middle-ear muscle reflex (MEMR) or the MOC reflex was activated, and whether such an activation could have affected the comparison between the attentional conditions.

In the analysed left ear, the stimulus waveforms were presented at 57 dB SPL, below the levels at which broadband noise typically elicits the MEMR [41, 42]. The speech in the right ear, at 69 dB SPL, lay well below the mean clinical MEMR threshold for broadband noise (about 81 dB SPL), but close to the mean threshold obtained with more sensitive wideband methods (about 68 dB SPL), both measured with the elicitor in the same ear as the reflex [43]. Since the reflex acts on both ears, an activation by the speech would also attenuate the transmission of sound through the left middle ear and thereby reduce the measured speech-like DPOAEs. Thresholds are, however, generally higher when the elicitor is presented to the opposite ear, as the speech was here, than when it is presented to the ear in which the reflex is measured [43, 44]. Moreover, even an activation of the reflex would have affected both attentional conditions alike, since the acoustic input was identical in both and the attended state of one voice was always recorded together with the ignored state of the other.Such an activation may have slightly reduced the strength of the emissions and thus the sensitivity to an attentional modulation, but it could not by itself have created differences between the conditions. The MOC reflex, in contrast, is activated by broadband sounds already at moderate levels; elicitors of about 60 dB SPL in the opposite ear are routinely used to evoke it [4, 39]. In the left ear, the reflex was therefore likely activated both by the stimulus waveforms in the same ear, at 57 dB SPL as in JS25, and by the speech in the right ear, at 69 dB SPL, about 12 dB louder than in JS25. For the same reason as above, this sound-evoked activation cannot by itself have created or masked differences between the attentional conditions. It does, however, set the state of the cochlea on which an attentional modulation would act: a stronger sound-evoked activation may have reduced the range left for an additional top-down modulation of cochlear gain. The higher level of the speech is therefore a further difference from JS25, besides the changes in stimulus design, that may have contributed to the absence of an attentional effect, and the present data cannot separate the two. The right ear, in which the speech was presented together with the stimulus waveforms, was not analysed here; the recordings ipsilateral to the speech will be reported separately.

### 4.6 Conclusion

Selective attention to one of two competing talkers did not robustly modulate speech-like DPOAEs in the present study, although the design was sensitive enough to detect an effect of the previously reported size. A simple model of the excitation patterns suggests a reason: with the two voices separated by only about one ERB, the emissions of the conditions most comparable to JS25 arose at cochlear places shared by the harmonics of both voices, where an attentional modulation would affect both voices alike. The isolated effects emerged only in conditions in which the shared places lay away from the stimuli and the emissions. Together with the strong effect in JS25, this suggests that an attentional modulation of speech-like DPOAEs, if it exists, depends on the separation of the competing voices and possibly on the bandwidth of the stimuli. The higher level of the speech, which may have activated the MOC reflex more strongly than in JS25, is a further difference that the present data cannot separate from the stimulus design. A follow-up study addressing these factors is planned.

## DATA AVAILABILITY

The data analysed in this study are available on Zenodo: https://doi.org/10.5281/zenodo.24104371.

## CODE AVAILABILITY

All custom code used for stimulus generation, analysis of speech-like DPOAEs and statistical evaluation is openly available on GitHub at https://github.com/janna-stb/speechlike_dpoae_followup under the MIT License.

## AUTHOR CONTRIBUTIONS

J.S.: Conceptualisation, Methodology, Software, Investigation, Formal analysis, Visualisation, Writing – original draft; T.R.: Conceptualisation, Supervision, Resources, Funding acquisition, Writing – review & editing.

## DECLARATION OF GENERATIVE AI AND AI-ASSISTED TECHNOLOGIES IN THE MANUSCRIPT PREPARATION PROCESS

During the preparation of this work, the authors used Claude (Anthropic) to formulate, revise and edit the text of the manuscript. In addition, Claude was used as a supporting tool during the development of the analysis code, for instance to revise, extend and debug existing code. All code was reviewed, tested and validated by the authors. After using this tool, the authors reviewed and edited the content as needed and take full responsibility for the content of the published article.

